# Cell-specific modulation of nuclear pore complexes controls cell cycle entry during asymmetric division

**DOI:** 10.1101/203232

**Authors:** Arun Kumar, Priyanka Sharma, Zhanna Shcheprova, Anne Daulny, Trinidad Sanmartín, Irene Matucci, Charlotta Funaya, Miguel Beato, Manuel Mendoza

## Abstract

The acquisition of cellular identity is coupled to changes in the nuclear periphery and nuclear pore complexes (NPCs). Whether and how these changes determine cell fate remains unclear. We have uncovered a mechanism regulating NPC acetylation to direct cell fate after asymmetric division in budding yeast. The lysine deacetylase Hos3 associates specifically with daughter cell NPCs during mitosis to delay cell cycle entry (Start). Hos3-dependent deacetylation of nuclear basket and central channel nucleoporins establishes daughter cell-specific nuclear accumulation of the transcriptional repressor Whi5 during anaphase and perinuclear silencing of the *CLN2* gene in the following G1 phase. Hos3-dependent coordination of both events restrains Start in daughter but not in mother cells. We propose that deacetylation modulates transport-dependent and - independent functions of NPCs, leading to differential cell cycle progression in mother and daughter cells. Similar mechanisms might regulate NPC functions in specific cell types and/or cell cycle stages in multicellular organisms.

## Introduction

Asymmetric cell division is a conserved mechanism that generates diversity in cell populations. Asymmetric divisions are present not only in unicellular organisms such as bacteria, budding yeasts, and flagellates, but also play a major role in stem cell self-renewal and tissue homeostasis during metazoan development (Knoblich, 2010; Li, 2013). During asymmetric division, unequal partitioning of cell fate determinants ultimately leads to distinct identities in the newly generated cells. This usually requires the establishment of a cell polarity axis that directs the distribution of fate determinants. Subsequent positioning of the cell division plane perpendicular to the fate determinant axis leads to their asymmetric partitioning between daughter cells after cytokinesis.

The nuclear envelope (NE) limits free diffusion of molecules between the nucleus and the cytoplasm. Transport between these two compartments is mediated by nuclear pore complexes (NPCs), macromolecular assemblies composed of approximately 30 nucleoporins that form a channel across the NE (Ibarra and Hetzer, 2015; Knockenhauer and Schwartz, 2016; Terry et al., 2007). Nucleo-cytoplasmic transport of proteins and RNA determines chromatin accessibility to transcription factors and mRNA export, and therefore is intimately tied to the regulation of gene expression and cell fate determination (Capelson et al., 2010a; Yang et al., 2013). In addition, the NE and nucleoporins associated with the nuclear basket of NPCs can directly interact with the nuclear genome to regulate gene expression and thus affect cell differentiation [reviewed in (Akhtar and Gasser, 2007; Capelson et al., 2010a; Mattout et al., 2015; Zuleger et al., 2014)]. In particular, the nuclear periphery is a transcriptionally repressive environment in yeast and metazoans (Andrulis et al., 1998; Green et al., 2012; Guelen et al., 2008; Pickersgill et al., 2006), and gene repositioning from the nuclear interior to the periphery can result in silencing (Kosak et al., 2002; Zink et al., 2004). Importantly, the composition of both the NE and NPCs, and the interactions of these elements with the genome, are known to diverge during development (D'Angelo et al., 2012; Korfali et al., 2014; Liang et al., 2013; Solovei et al., 2013), further suggesting that modifications at the nuclear periphery might play a role in the acquisition or maintenance of cell identity. However, how differences in perinuclear function are established during development, and in particular during asymmetric cell divisions, remains unclear.

Budding yeast divide asymmetrically, giving rise to mother and daughter cells of different size, age, transcriptional profiles and cell cycle programs (Colman-Lerner et al., 2001; Hartwell and Unger, 1977; Shcheprova et al., 2008). In particular, commitment to a new division cycle is regulated asymmetrically in *S. cerevisiae*: daughter cells start a new cycle later than mother cells. This is due both to a cell size-dependent delay that prolongs G1 until daughter cells reach a critical size (Hartwell and Unger, 1977; Turner et al., 2012) and a size-independent, daughter-specific delay of the G1/S transition (Di Talia et al., 2009; Laabs et al., 2003).

The regulatory principles controlling cell cycle entry are similar in yeast and animal cells. In both cases, a transcriptional activator (SBF in yeast; E2F in mammals) drives expression of cyclin genes (*CLN1/2* or cyclin E, respectively) controlling the start of S phase. SBF or E2F are inhibited in G1 by a transcriptional repressor: Whi5 in yeast, and its homolog the Rb tumour suppressor in mammals. In yeast, a key event driving the G1/S transition is the dilution of Whi5 activity by cell growth, whereby the volume increase in daughter cells during G1 lowers the concentration of Whi5 below a critical threshold (Schmoller et al., 2015). This allows Cyclin-dependent kinase (Cdk) complexes to inactivate Whi5, which is then evicted from the nucleus (Costanzo et al., 2004; de Bruin et al., 2004). Interestingly, the G1 concentration of Whi5 is higher in daughter cells than in mother cells (Liu et al., 2015; Schmoller et al., 2015). The mechanism establishing this Whi5 concentration asymmetry is not known.

Here, we discovered that the distinct cell cycle program in budding yeast daughters compared to mothers is due to daughter-specific association of the class II histone deacetylase (HDAC) Hos3 with NPCs. We identify the mechanism that recruits Hos3 to the NPCs in the daughter cell during mitosis. Further, we demonstrate that Hos3-mediated lysine deacetylation of distinct nucleoporins within NPCs establishes asymmetric segregation of the Whi5 transcriptional repressor and perinuclear tethering of the G1/S cyclin gene *CLN2* in daughters. We find that both of these Hos3-coordinated functions contribute to inhibit Start in daughter cells. Thus, cell-specific regulation of NPCs by lysine deacetylation can direct differences in cell identity during asymmetric division.

## Results

### Hos3 inhibits cell cycle entry exclusively in daughter cells

Commitment to a new division cycle in budding yeast occurs first in mother cells, and only later in daughter cells. Although the lysine deacetylase Hos3 has been implicated in the control of G1 length and gene expression, it is not known to control the distinct cell cycle programs in mother and daughter cells (Huang et al., 2009; Wang et al., 2009). To investigate this hypothesis, we determined G1 duration in wild type and *hos3*Δ mother and daughter cells by evaluating the interval between cytokinesis and bud emergence. Specifically, we measured the disappearance of Myo1-GFP from the bud neck to its reappearance to the new bud site, normalized by the rate of growth α during this period (αTG1) (Figure **1A**). This revealed a shortened G1 in *hos3*Δ daughter cells, but not mother cells, relative to wild type (Figure **1B-C**, **S1A** and Supplementary **Table S1**).

**Figure 1:**
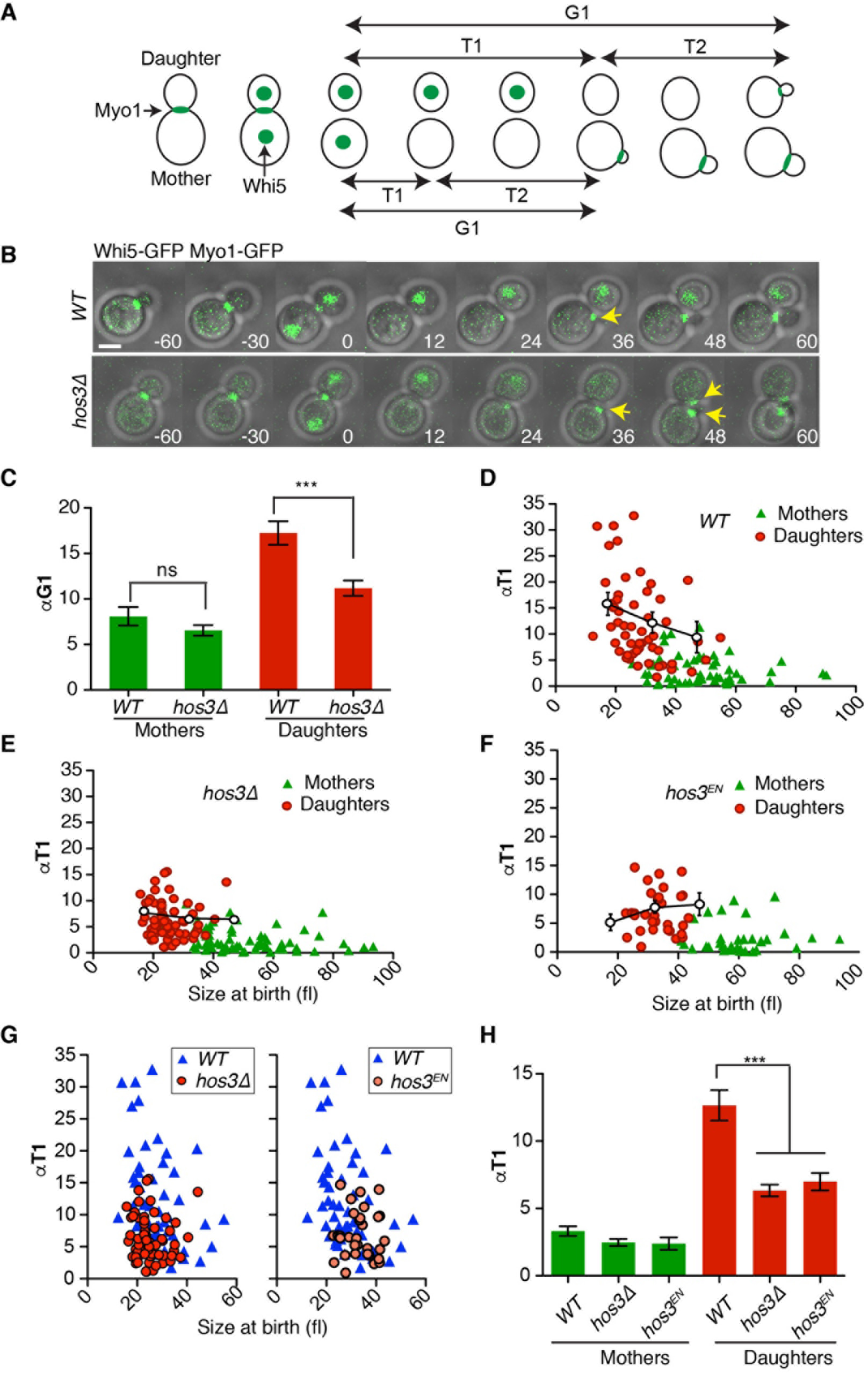
Hos3 delays Start in daughter cells. (**A**) Scheme showing the separation of G1 in two distinct periods, T1 and T2, relative to Myo1 and Whi5 dynamics. **(B)** Composite of phase contrast, Whi5-GFP and Myo1-GFP in wild type (*WT*) and *hos3*Δ cells. Daughter *hos3*Δ cells start a new bud (Myo1-GFP appearance, arrows) earlier than *WT*. Confocal sections spanning the entire cell were acquired at 3-minute intervals. Time is indicated in minutes; t=0 marks the last frame before cytokinesis. Scale bars, 2 μm. (**C**) αTG1 values (mean and SEM) and (**D-G**) correlation between αT1 (with binned means and SEM) and cell size at the time of birth (cytokinesis), for mother and daughter cell pairs in *WT* (110 cells), *hos3*Δ (128 cells) and *hos3*^*EN*^ (66 cells). **(H)** αT1 values (mean and SEM) are shown in cells of the indicated strains (***, *p* < 0.001; *p* > 0.05; Mann-Whitney test).

G1 is divided in two periods, T1 and T2, separated by nuclear export of the transcriptional repressor Whi5 (Figure **1A**). Whi5 nuclear export determines commitment to the next cell cycle (Start) by allowing expression of MBF/SBF target genes required for budding and DNA replication (Turner et al., 2012). Analysis of Whi5-GFP nuclear export kinetics (T1) showed that *HOS3* deletion reduces αT1 in daughter cells relative to wild-type daughters (mean and standard deviation: *WT*: 12.6 ± 8.3, N = 55; *hos3*Δ: 6.3 ± 3.5, N = 64, *p* < 0.0001, Mann-Whitney test) (Figure **1, D-E**). Shortening of G1 in *hos3*Δ daughter cells relative to wild type daughters was observed even for cells born at similar sizes (Figure **1G**). In contrast, the cellular growth rate and the scaled time between Whi5 export and budding (αT_2_), ascribed to molecular noise (Di Talia et al., 2007), were similar in *hos3*Δ and wild type cells (Figure **S1B-C**). In addition, a catalytically inactive mutant of Hos3 (Hos3^H196E, D231N^, or Hos3^EN^) (Wang and Collins, 2014) showed advanced cell cycle entry exclusively in daughter cells in a similar manner to *hos3*Δ (Figure **1F-H**). Therefore, Hos3 activity inhibits Start in daughter cells.

### Hos3 is recruited to the nuclear periphery of daughter cells during anaphase

Hos3 is present in the cytoplasm during interphase but is recruited to the daughter side of the septin-based ring at the mother-bud neck during mitosis (Wang and Collins, 2014). To further investigate the subcellular localization of Hos3-GFP during mitosis, we performed time-lapse fluorescence microscopy. Unexpectedly, we discovered that Hos3-GFP associates with the nuclear periphery of the daughter cell, but not the mother cell, during anaphase. The perinuclear localization of Hos3 in daughter cells is coincident with migration of the anaphase nucleus from mother to daughter through the bud neck (Figure **2A**, **S2A**, movies **S1-2**). Hos3 disappeared from both the bud neck and the nuclear periphery 1-2 minutes before cytokinesis, marked by contraction of the acto-myosin ring and septin ring splitting, visualized by mCherry-tagged Myo1 and Cdc3, respectively (Figure **S2B-C** and movie **S3**). Hos3 protein levels did not change dramatically during mitosis, suggesting that protein degradation is not the major driver of its localization dynamics (Figure **1B** and **S2D**). The catalytically inactive protein Hos3^EN^-GFP showed enrichment at the bud neck and nuclear periphery identical to the wild type protein (Figure **S2E**), indicating that the deacetylase activity of Hos3 is not required for its localization dynamics.

**Figure 2:**
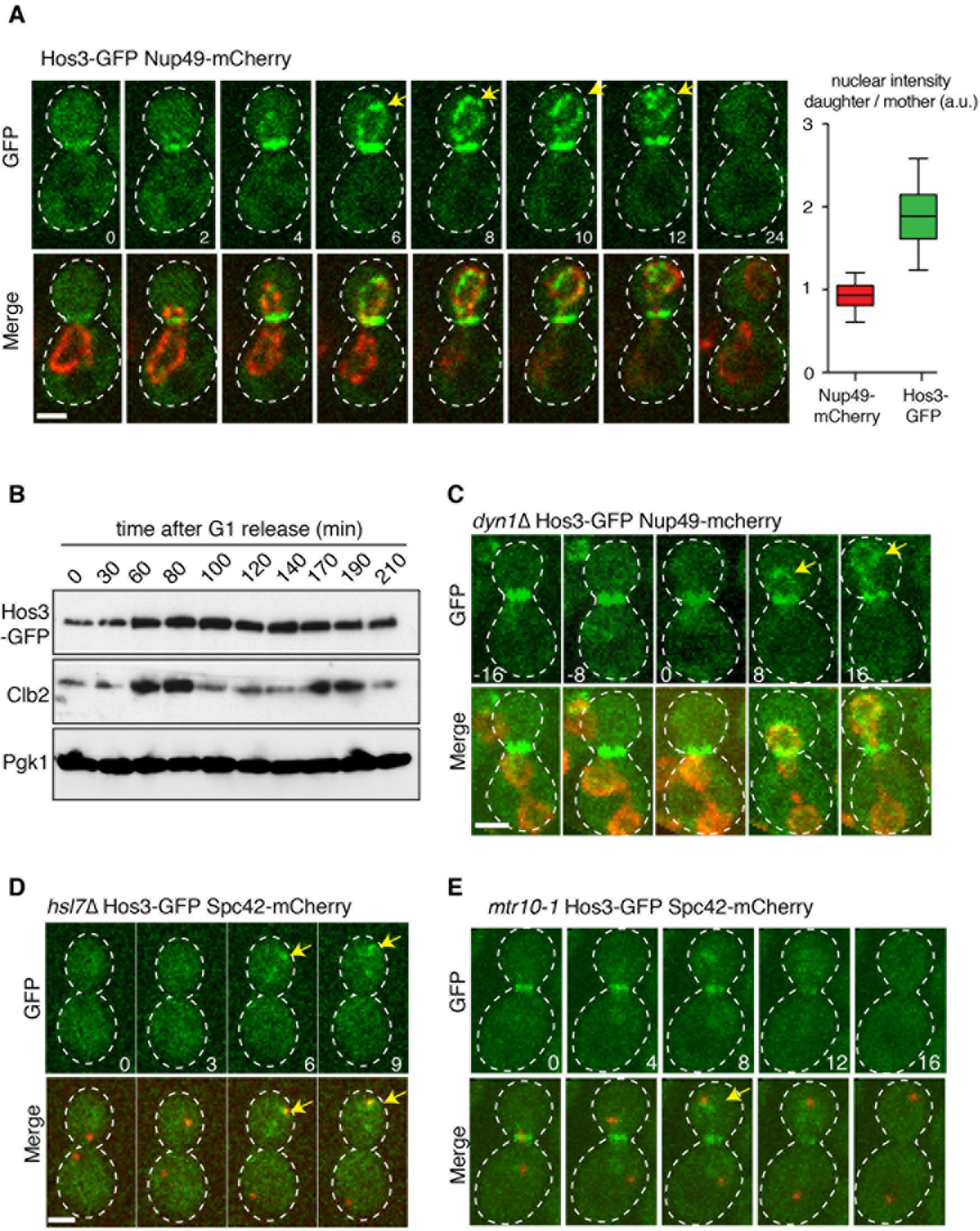
Hos3 localizes to the periphery of daughter nuclei upon nuclear migration in anaphase. **(A)** Localization of Hos3-GFP (green) to the bud neck during mitosis and to the daughter nuclear periphery (Nup49-mCherry, in red) during nuclear migration into the bud. Arrows point to perinuclear Hos3-GFP. The graph shows the ratio of nuclear intensity of GFP and mCherry signals in late anaphase cells (N = 67). **(B)** Cell extracts were prepared at the indicated times after release from a G1 block (induced with alpha factor). Hos3-GFP was detected with an anti-GFP antibody; Clb2 (B-type cyclin) serves as a cell cycle progression marker, and Pgk1(3-phosphoglycerate kinase) as loading control. (**C**) A *dyn1*Δ cell showing separated anaphase nuclei (Nup49-mCherry, in red) in the mother cell. Hos3-GFP associates with the nuclear periphery only after nuclear migration across the bud neck (arrows). (**D**) Hos3-GFP is absent from the bud neck and nuclear periphery in the *hsl7*Δ mutant, but localizes to the dSPB (marked with Spc42-mCherry, red) in anaphase (arrows). **(E)** Hos3-GFP localizes to bud neck and spindle pole body (marked with Spc42-mCherry) in *mtr10-1* cells at 30°C, but fails to localize at the nuclear periphery. Maximal projections of whole-cell Z-stacks are shown, expect in (A) where one medial section is shown for clarity. Images were acquired at 2-4 minute intervals; time is indicated in minutes. t=0 is the last frame before the nucleus traverses the bud neck. Scale bars, 2 μm.

We next asked whether Hos3 enrichment at the nuclear periphery is dependent on passage of the nucleus across the bud neck and into the daughter cell. To test this hypothesis, we first examined Hos3-GFP localization in cells arrested in metaphase by depletion of the APC activator Cdc20; these cells develop nuclear protrusions extending across the bud neck and into the daughter cell compartment (Kirchenbauer and Liakopoulos, 2013). Hos3-GFP decorated nuclear protrusions and remained confined in the daughter cell (Figure **S3A**). Next, we examined cells that completed anaphase within the mother cell due to loss of the dynein microtubule motor Dyn1. In *dyn1*Δ mutants, Hos3-GFP was recruited to the perinuclear region only after migration of the anaphase nucleus into the daughter compartment (20/20 cells with delayed nuclear migration) (Figure **2C**, **S3B-C**, and movie **S4**). Thus, nuclear migration through the bud neck is required for Hos3 targeting to the perinuclear region.

We next investigated the relationship between the bud neck localization of Hos3 in early mitosis and its subsequent perinuclear enrichment. Hos3 localization to the bud neck depends on septins and the septin-interacting protein, Hsl7 (Wang and Collins, 2014). We found that mild perturbation of septin ring structure in temperature sensitive (ts) *cdc12-1* mutant cells led to symmetric localization of Hos3 at both the bud neck and the nuclear periphery during anaphase (Figure **S4**). Moreover, deletion of Hsl7 prevented perinuclear localization of Hos3-GFP, which was instead detected at the anaphase daughter spindle pole body (dSPB) (71/74 cells) (Figure **2D** and **S5A**). Thus, proper recruitment of Hos3 to the septin ring is required for its enrichment at the nuclear periphery of daughter cells.

We note that enrichment of Hos3 at the dSPB, but not at the nuclear periphery, was reported previously using ectopically expressed Hos3-GFP (Wang and Collins, 2014). Indeed, we observed a similar localization pattern when Hos3-GFP was expressed from the strong *ADH1* promoter (Figure **S5B**, movie **S5**), suggesting that Hos3 is preferentially detected at the dSPB when over-expressed. Hos3 was also enriched at the dSPB after inhibition of mitotic exit in *cdc15-1* ts mutants at 37 °C, which in addition retained Hos3-GFP in the bud neck (Figure **S5C**). We conclude that Hos3 recruitment to the bud neck is dispensable for its localization to the dSPB but is essential for its enrichment at the daughter cell nuclear periphery.

Finally, we asked whether the perinuclear localization of Hos3 reflects an association with the cytoplasmic or the nucleoplasmic side of the nuclear envelope. Inactivation of nuclear import factors Kap95, Kap60 and Kap122 did not affect the perinuclear localization of Hos3-GFP (data not shown). In contrast, inactivation of the importin Mtr10 impaired Hos3 recruitment to the nuclear periphery, which was instead found at the bud neck and the dSPB during anaphase (42/52 cells; Figure **2E**). Notably, changes in Hos3-GFP localization in *dyn1*, *hsl7* and *mtr10* cells were not due to differences in Hos3-GFP levels, as these were comparable across the mentioned strains (Figure **S5D**). Together, the above data suggest that Hos3 is imported into the daughter nucleus during its migration across the bud neck, in a manner dependent on Mtr10. The highly regulated asymmetric enrichment of Hos3 at the nuclear periphery in daughters suggests that it performs an important function in daughter cell nuclei, and might underlie the Hos3-mediated inhibition of Start in daughters.

### Hos3 associates with nuclear pores in daughter cells to delay Start

To examine whether perinuclear Hos3 is sufficient to delay Start, we asked if targeting Hos3 to the mother cell nucleus prolongs G1 in these cells. We generated Hos3-GFP fused with the nuclear localization signal of SV40 TAg (Kalderon et al., 1984) (Hos3-NLS-GFP) and found that it localized to the periphery of both mother and daughter cell nuclei throughout the cell cycle (Figure **3A** and **S6A-B**). Importantly, we found that Whi5 nuclear export (T1) was delayed in both mother and daughter cells expressing symmetrically distributed Hos3-NLS by time-lapse imaging (Figure **3**, **B-C**). Furthermore, over-expression of Hos3-NLS impaired growth on solid media (Figure **3D** and **S6C**) and led to a dramatic increase in cell volume, consistent with a severe cell cycle delay (Figure **3E**). These phenotypes arising from Hos3-NLS over-expression required both the constitutive nuclear localization and the catalytic activity of Hos3, as cells grew normally when over-expressing Hos3-GFP (without the NLS) or Hos3^EN^-NLS-GFP (Figure **3D** and **S6C**). We conclude that the perinuclear localization of active Hos3 is sufficient to inhibit Start.

**Figure 3:**
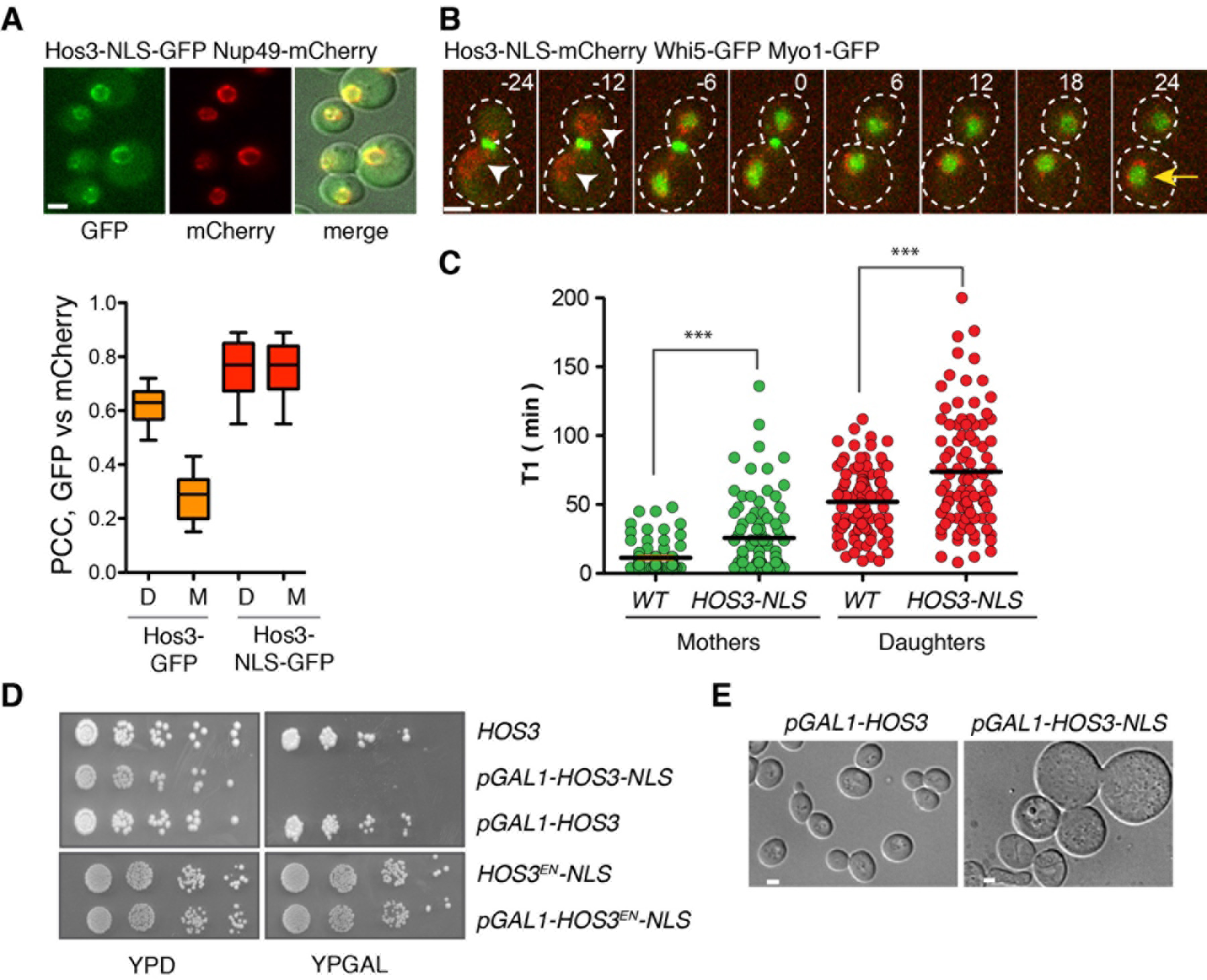
Perinuclear Hos3 inhibits cell cycle entry. (**A**) Hos3-NLS-GFP (green) localizes to the nuclear periphery (Nup49-mCherry, red) in mother and daughter cells. The graph shows the Pearson's correlation coefficient of the GFP and mCherry channels in the indicated anaphase cells (N > 28). (**B**) Anaphase and G1 in a cell expressing Hos3-NLS-mCherry. Arrowheads point to perinuclear Hos3-NLS in mother and daughter cells; the arrow points to persistent Whi5-GFP in the mother nucleus. Time is indicated in minutes; t=0 marks the last frame before cytokinesis. (**C**) Duration of pre-Start G1 phase (T1) in *WHI5-GFPMYO1-GFP* cells, expressing either Hos3 (*WT*) or Hos3-NLS-mCherry (> 174 cells per strain). Lines represent the mean. ***, *p* < 0.001, Mann-Whitney test. (**D**) Growth inhibition upon over-expression of Hos3-NLS from the *GAL1,10* promoter. Serial dilutions of the indicated strains expressing Hos3-GFP fusions were incubated for 2 days at 30 °C. (**E**) DIC images of wild type and Hos3-NLS after 2 days of growth in YP + galactose plates. Scale bars, 2 μm.

Notably, Hos3-NLS did not accumulate in the nucleoplasm but was instead enriched in the nuclear periphery like its wild-type counterpart (see Figure **3A**), and was detected by immuno-electron microscopy on the nucleoplasmic side of nuclear pore complexes (NPCs) (Figure **S6D**). Moreover, artificial NPC clustering caused by deletion of *NUP133* (Pemberton et al., 1995) led to clustering of Hos3-GFP and Hos3-GFP-NLS, which partially co-localized with Nup60-mCherry (Figure **4A**, **S7A-C**, and movie **S6**). These findings suggested that Hos3 associates with NPCs. To test this hypothesis, we investigated the role of distinct NPC components in the perinuclear localization of Hos3. We found that the perinuclear association of Hos3-NLS was independent of the nuclear pore basket components Nup2 and Mlp1/2, but was dependent on Nup60 (Rout et al., 2000) (Figure **S7D-E**). By time-lapse microscopy, we found that although Hos3-GFP localized to the anaphase daughter nuclei in *nup60*Δ cells, it was no longer retained at the nuclear periphery. Instead, Hos3-GFP was homogeneously distributed throughout the nucleoplasm in *nup60*Δ anaphase cells (Figure **4B** and movie **S7**). Furthermore, we detected a direct interaction between epitope-tagged Hos3 and Nup60 in logarithmically growing cultures by co-immunoprecipitation. Interestingly, this interaction was much stronger in extracts prepared from Cdc20-depleted cells that are arrested in metaphase, when Hos3 localizes to the nuclear periphery (Figure **4C** and **S7F**), supporting the notion that Hos3 associates with nuclear pore basket components during nuclear migration into the bud.

**Figure 4:**
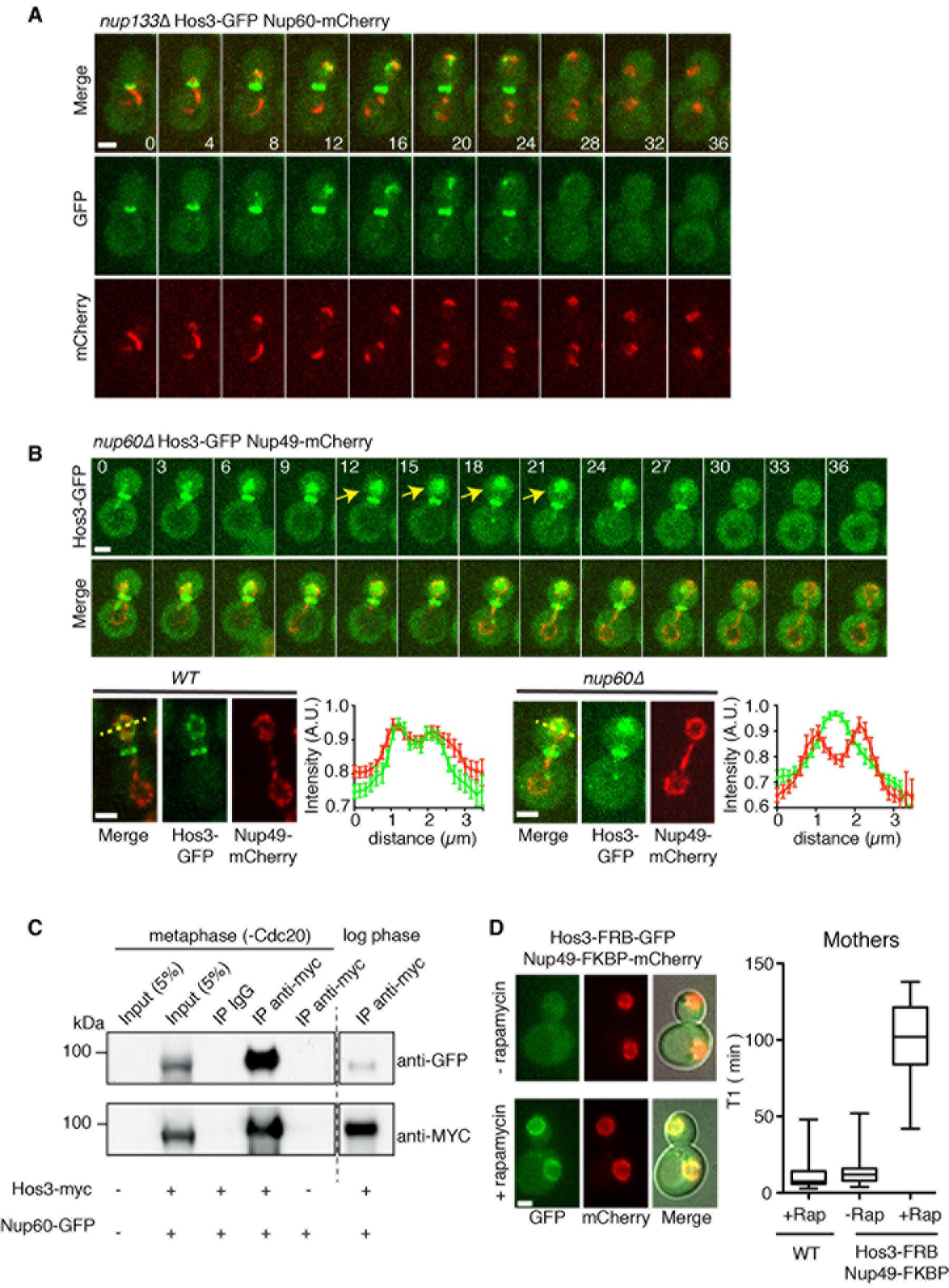
Hos3 associates with nuclear pore complexes. **(A)** Hos3-GFP (green) Nup60-mCherry (red) form partially overlapping clusters during anaphase in *nup733*Δ cells. (**B**) Loss of perinuclear Hos3-GFP and its distribution in the nucleoplasm during anaphase in a *nup60*Δ cell. The fluorescence intensity of Hos3-GFP and Nup49-mCherry (to mark the nuclear periphery) were measured across the daughter cell nucleus in 10 wild type and 10 *nup60*Δ late anaphase cells (yellow lines). Hos3-GFP and Nup49-mCherry profiles closely overlap in wild type whereas Hos3 is distributed throughout the nucleoplasm in *nup60*Δ. **(C)** Hos3 and Nup60 are associated in mitotic yeast extracts. Hos3-myc was immunoprecipitated in extracts prepared from metaphase-arrested cells (Cdc20 depletion) and from asynchronously growing cultures (log phase). Proteins were detected by western blot using the indicated antibodies. **(D)** Hos3-FRB-GFP recruitment to NPCs (co-localization with Nup49-FKBP-RFP) upon rapamycin addition prolongs T1 duration in mother cells (right; N=25 cells). Rapamycin (25 μM) was added immediately before imaging. Association of Hos3-GFP-FRB with nuclear pores was observed within 5 minutes of rapamycin addition. T1 in daughter cells was inhibited for the duration of the movie (3 hours) and is not plotted. Lines represent the mean. In (A-B), time is indicated in minutes; t=0 marks the last frame before nuclear entry into the bud. Scale bars, 2 μm.

To test if Hos3 interactions with the NPC could play a role in the inhibition of Start, we determined if artificial tethering of Hos3 to NPCs with an FKBP-FRB system (Gallego et al., 2013) affected the time from cytokinesis to Whi5 nuclear export. Indeed, we found that addition of rapamycin to *HOS3-FKBP NUP49-FRB* cells caused immediate recruitment of Hos3 to the nuclear periphery, and strongly and specifically delayed Start (Figure **4D**). We conclude that retention of Hos3 at the nuclear periphery through interactions with the NPC during anaphase is sufficient to delay Whi5 export and cell cycle entry.

### Hos3 is required for the enrichment of Whi5 in daughter cell nuclei

To gain insight into the mechanism by which Hos3 inhibits Start in daughter nuclei, we next asked whether the presence of Hos3 alters the protein composition at the nuclear periphery. To facilitate the identification of proteins whose subnuclear distribution is regulated by Hos3, we examined the localization of 217 nuclear proteins fused to GFP, in wild type and *HOS3-NLS* cells (Figure **5A-B** and Supplementary **Table S2**). We did not observe Hos3-dependent changes for 207 of these candidates, including more than 30 nucleoporins.

**Figure 5:**
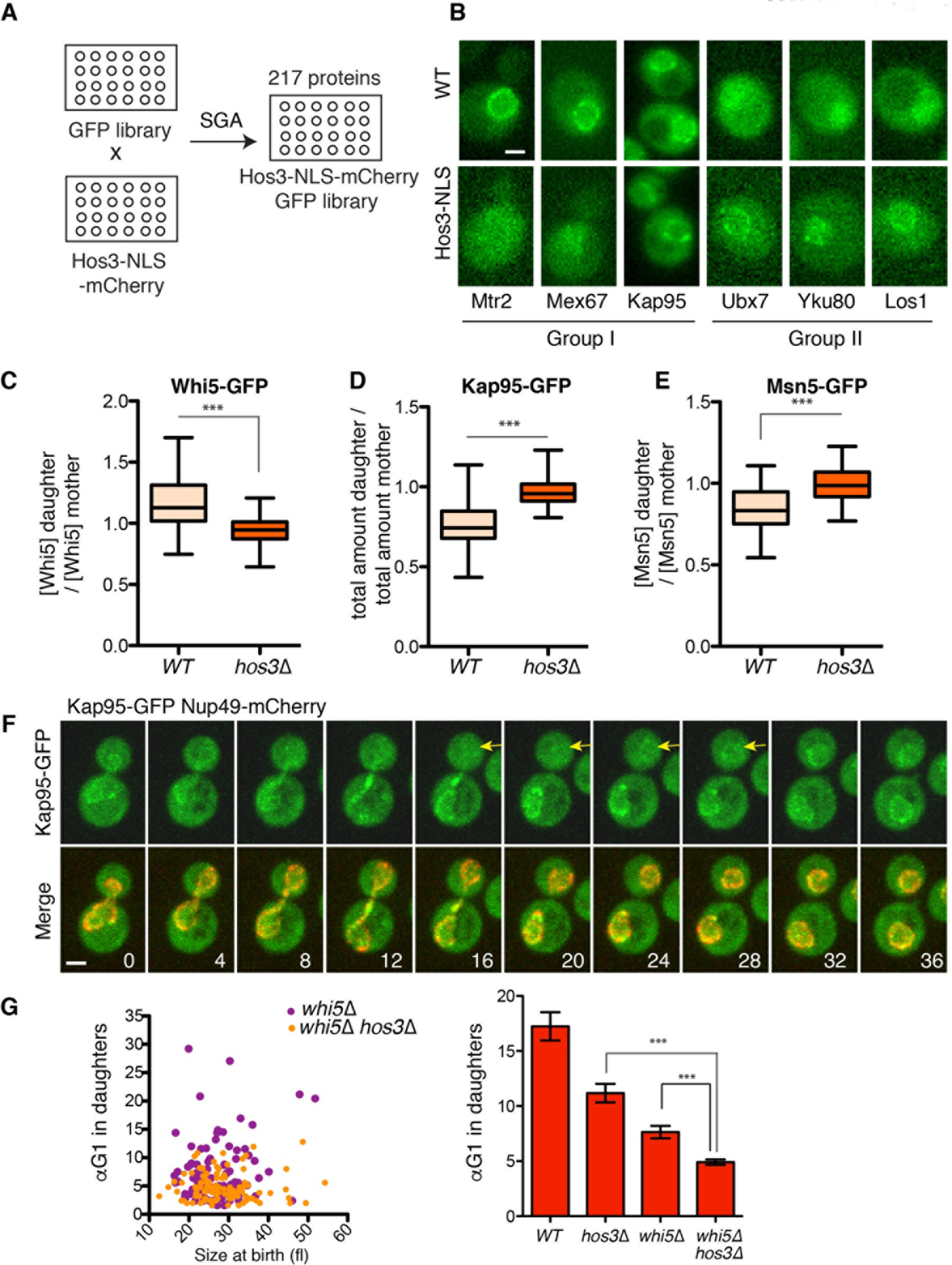
Hos3 modulates localization of perinuclear factors and controls the asymmetry of Whi5 and karyopherins. **(A)** Construction of a library of nuclear GFP-fusion proteins expressing Hos3 or Hos3-NLS. **(B)** Examples of protein localization changes induced by Hos3-NLS. **(C-E)** Hos3 affects asymmetric distribution of Whi5, Kap95 and Msn5. Nuclear fluorescence was measured in mother and daughter cells at the time of birth (> 50 cells / strain; ***, *p* < 0.001, Mann-Whitney test). **(F)** Localization of Kap95-GFP (green) at nuclear periphery (marked with Nup49-mCherry, in red) in mother and daughter cells during cell division. Arrows point to low perinuclear Kap95-GFP in daughter cells. **(G)** Correlation between αTG1 and cell size at the time of birth (cytokinesis), for mother and daughter cell pairs in *whi5*Δ (178 cells) and *whi5*Δ *hos3*Δ (204 cells) (*** *p* < 0.001, Mann-Whitney test). Strains from Figure 2 are included for comparison. Note that TG1 was used to measure Start instead of T1, since the latter is defined by Whi5 nuclear exit. Scale bar, 2 μm.

In contrast, 10 proteins showed altered localization in *HOS3-NLS* relative to wild type control strains, and belonged to two major categories. Group I includes proteins that showed reduced perinuclear localization in Hos3-NLS cells. These were all involved in nucleo-cytoplasmic transport and comprised the karyopherins Kap95 and Kap123, the mRNA export regulators Mtr2 and Mex67, and the TREX-2 component Sac3 (Figure **5B** and **S8A**). Group II proteins showed the inverse pattern: these proteins localized to the nucleoplasm in wild type cells and became enriched at the nuclear periphery in the presence of Hos3-NLS. This group was more heterogeneous and included the tRNA export factor Los1, the telomeric Ku complex components Yku70 and Yku80, and the UBX domain-containing Ubx7 (Figure **5B** and **S8A**). In addition, the karyopherin Kap114 was weakly enriched in the nucleoplasm of wild type cells, and this enrichment was increased in Hos3-NLS cells.

The identification of group I proteins as potential Hos3 effectors raised the possibility that Hos3 regulates nucleo-cytoplasmic transport dynamics to delay Start in daughter cells. The nuclear concentration of the S-phase inhibitor Whi5 is intimately linked to the regulation of the G1/S transition. Indeed, the probability of entering the cell cycle is inversely proportional to the Whi5 nuclear concentration, and Start is triggered when cell growth dilutes Whi5 below a critical level (Liu et al., 2015; Schmoller et al., 2015). Notably, the Whi5 nuclear concentration is higher in daughter than mother cells, although the reason for this asymmetry has not been elucidated. We therefore determined the nuclear concentration of Whi5-GFP in wild type and *hos3Δ* cells right after cytokinesis. We found that the Whi5 nuclear concentration was higher in daughter than in mother cells. This asymmetry was slightly lower than previously described, possibly owing to differences in strain background or imaging conditions. Strikingly however, the asymmetric nuclear concentration of Whi5 was dependent on Hos3 (Figure **5C** and **S8B**).

We next focused on the regulation of karyopherins by Hos3. Specifically, we asked whether Hos3 regulates the nuclear levels of the Kap95 importin and of the Msn5 exportin, which together control Whi5 nuclear levels (Kosugi et al., 2009; Taberner et al., 2009). We found that after cytokinesis, the total amount and the concentration of Kap95-GFP and Msn5-GFP are higher in mother than in daughter nuclei. Furthermore, these asymmetries were dependent on Hos3 (Figure **5D-F** and **S8C-D**). Thus, nuclear Kap95 and Msn5 are distributed asymmetrically and with the opposite polarity of their cargo Whi5, which is more concentrated in daughter than in mother nuclei. Notably, deletion of Hos3 did not affect the asymmetric distribution of NPCs marked with Nup49-mCherry (Figure **S8E**), which are partially retained in mother cells due to a septin-based diffusion barrier (Shcheprova et al., 2008). These results suggest that Hos3 inhibits cell cycle entry in daughter cells through modulation of Whi5 nucleo-cytoplasmic transport in late anaphase. We propose that this leads to a higher Whi5 concentration in daughter nuclei, and consequently prolongs G1 duration of these cells.

To test whether regulation of Whi5 levels accounts entirely for the role of Hos3 in Start, we examined G1 length in cells lacking both Hos3 and Whi5. Deletion of *WHI5* leads to advanced Start due to derepression of SBF/MBF target genes (Costanzo et al., 2004; de Bruin et al., 2004). Intriguingly, *hos3*Δ *whi5*Δ cells showed shorter G1 duration than either single mutant (Figure **5G** and **S9**). This suggests that in addition to its role controlling Whi5 nuclear levels, Hos3 delays Start through additional, Whi5-independent mechanism(s).

### Hos3 inhibits *CLN2* expression in parallel to the *CLN3* repressor Ace2

To further investigate how the Hos3 interaction with the NPC during anaphase delays Start in daughter cells, we examined the relationship between Hos3 and known regulators of cell cycle entry. Control of Start is composed of two distinct modules, each controlled by a different G1 cyclin (Di Talia et al., 2007). Cln3-Cdk complexes initiate Start through inactivation of Whi5 in early G1, followed by Cln1/2-Cdk complexes, which complete Whi5 inactivation in a positive feedback loop (Cross and Tinkelenberg, 1991; Dirick and Nasmyth, 1991; Skotheim et al., 2008) (Figure **S10A**). The daughter-specific factor Ace2 delays G1 in daughters by repressing *CLN3* (Di Talia et al., 2009; Laabs et al., 2003). As Ace2 is retained in daughter nuclei during G1, we asked whether its localization dynamics depended on Hos3. However, the nuclear residence time of Ace2-GFP was not affected in *hos3*Δ cells (Figure **S10B**). Moreover, advanced Start in *hos3*Δ cells was independent of the presence of *CLN3*, but required the presence of *CLN2* (Figure **S10C-D**). Finally, deletion of Hos3 and Ace2 advanced Start to similar extents, and *hos3*Δ *ace2*Δ double mutants showed additive effects, suggesting that Ace2 and Hos3 function in distinct pathways (Figure **S10D-E**). Together, these results show that Hos3 delays Start in daughter cells in parallel to the Ace2 - Cln3 module, and possibly by inhibiting the Cln2-associated module.

### Hos3 controls *CLN2* expression and subnuclear localization of the *CLN2* gene locus

We next analyzed *CLN2* expression by time-lapse microscopy of wild type and *hos3*Δ cells expressing fast-folding, unstable GFP expressed from the *CLN2* locus under the control of the *CLN2* promoter (Bean et al., 2006). This analysis showed that the time of *CLN2* expression relative to cytokinesis, determined using Myo1-mCherry, was advanced in *hos3*Δ daughter cells (Figure **6A-B** and **S11**). The deacetylase Rpd3 binds to the *CLN2* promoter to repress it in G1 phase (Takahata et al., 2009; Wang et al., 2009). We could detect Rpd3 but not Hos3 at the *CLN2* promoter (Figure **S12**), suggesting that Rpd3 and Hos3 repress *CLN2* through different mechanisms.

**Figure 6:**
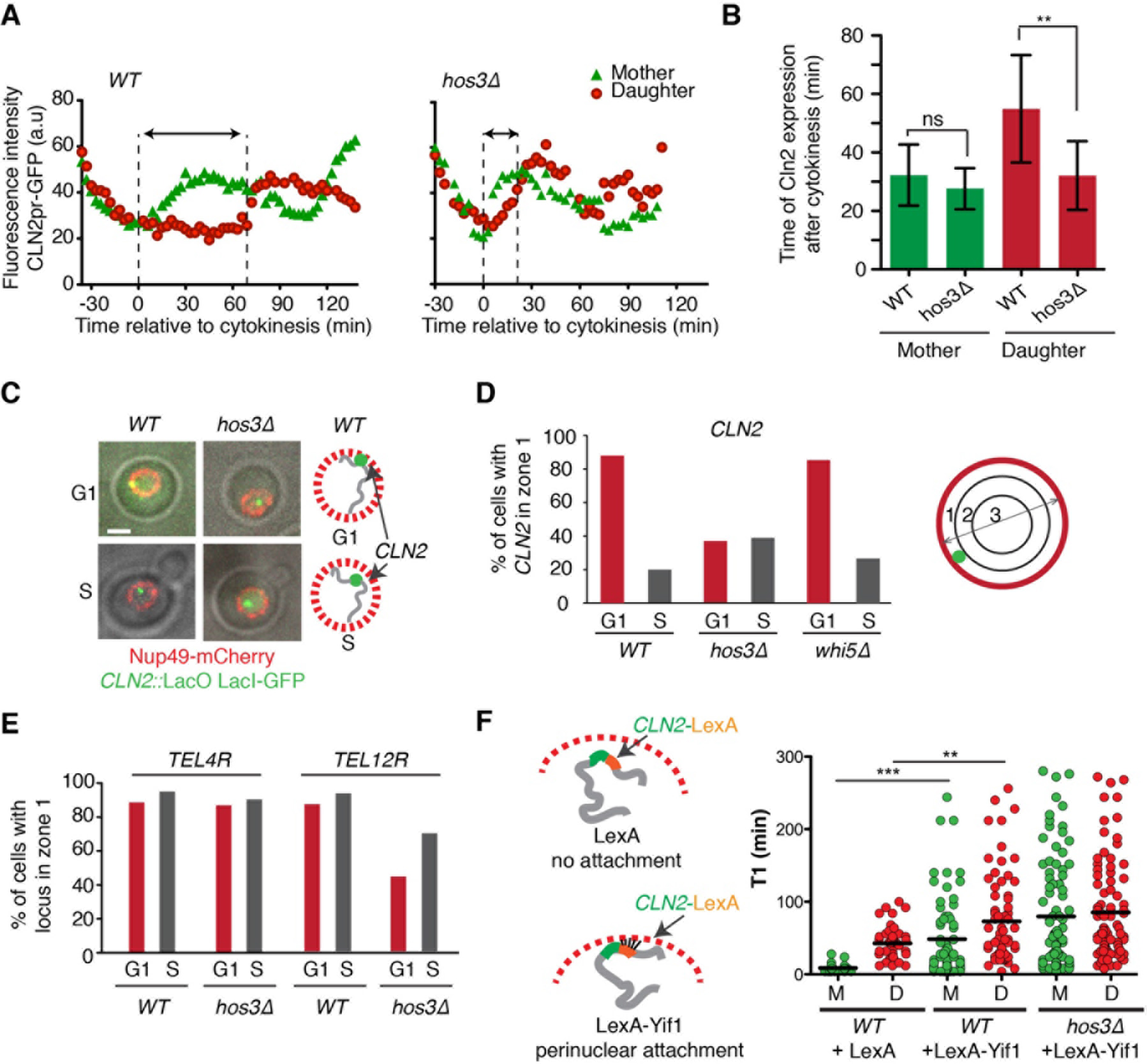
Hos3-dependent perinuclear anchoring of *CLN2* delays Start. **(A)** Fluorescence intensity of CLN2pr-GFP in mother and daughter cells relative to cytokinesis (Myo1-mCherry disappearance; t=0) in wild type and *hos3*Δ. Images were acquired at 3 min intervals. The GFP signal increases shortly after cytokinesis in *WT* and *hos3*Δ mother cells. Arrows indicate the interval between cytokinesis and the half-maximum of CLN2pr-GFP intensity in daughter cells. GFP accumulates in the wild type daughter cell only after a delay; this delay is shorter in the *hos3*Δ daughter cell. **(B)** Time (mean and SD) of half-maximum CLN2pr-GFP intensity relative to cytokinesis for *WT* (20 mother-bud pairs) and *hos3*Δ (15 pairs) (**, *p* < 0.01; ns, non-significant, *p* > 0.05, Student’s t-test). **(C)** Composite of phase contrast, *CLN2::lacO* (LacI-GFP, green) and Nup49-mCherry (red) in G1 and S-phase cells of the indicated strains. **(D-E)** Subnuclear position of *CLN2* **(D)** and two subtelomeric loci **(E)** in the indicated strains and cell cycle stages, determined by mapping their localization to one of three concentric nuclear zones of equal surface in confocal optical slices (N > 100 cells). **(F)** Duration of pre-Start G1 phase (T1) in cells of the indicated strains, in the absence and presence of artificial tethering of *CLN2* to the nuclear periphery (N > 92 cells per strain). Lines represent the mean (** *p* < 0.01, *** *p* < 0.001Mann-Whitney test).

The yeast nuclear periphery and NPCs play an important role in three-dimensional genome organization and gene expression. For instance, yeast telomeres are anchored at the nuclear periphery, which is associated with gene silencing (Taddei and Gasser, 2012; Zimmer and Fabre, 2011). As the *CLN2* gene locus is positioned near a telomere, just 65 kb away from *TEL16R* (Figure **S13A**), we hypothesized that *CLN2* might be positioned in the perinuclear region, where it would be poised for Hos3-dependent regulation. Confocal 3D imaging showed that *CLN2* labeled with the LacO/LacI-GFP system was adjacent to the NE (Nup49-mCherry) in 88% of unbudded cells and in 20% of budded cells (Figure **6C-D**, and **S13B-C**). Thus, *CLN2* is perinuclear in G1 and is displaced towards their nuclear interior in S phase. This cell-cycle dependent localization was specific to *CLN2*, because subtelomeric loci (*TEL4R* and *TEL12R)* localized to the NE region in a similar fraction (≈88%) of G1 and S-phase cells (Figure **6E**). Strikingly, the G1-specific perinuclear association of *CLN2* was independent of *WHI5* but required Hos3, as the fraction of G1 cells with *CLN2* at the NE was reduced to less than 40% in *hos3*Δ cells (Figure **6D** and **S13C**). Hos3 deletion also led to displacement of *TEL12R* but not of *TEL4R* from the NE, suggesting that Hos3 affects the perinuclear localization of multiple gene loci (Figure **6E** and **S13D**).

To determine if Hos3-dependent perinuclear association of *CLN2* might contribute to the role of Hos3 in Start, we evaluated ectopic recruitment of *CLN2* to the NE. *CLN2* containing LexA recognition sites within the 3’UTR was recruited to the NE via a LexA-Yif1 fusion protein (Schober et al., 2009). Expression of LexA-Yif1, but not of LexA alone, increased G1 duration in both mother and daughter cells, indicating that targeting of *CLN2* to the NE is sufficient to delay Start. This delay was not bypassed in *hos3*Δ cells, suggesting that Hos3 targets *CLN2* to the nuclear envelope but is dispensable for its repression once it is localized at the NE (Figure **6F**).

### Daughter cell cycle control requires deacetylation of NPC components

The association of Hos3 with Nup60 raised the possibility that at least some of its functions in cell cycle entry might involve its association and/or deacetylation of NPC components. Consistent with the first possibility, time-lapse imaging showed that deletion of *NUP60* impaired *CLN2* perinuclear recruitment, but did not affect Whi5 asymmetric segregation and only moderately reduced G1 length in daughters (Fig. **S14**). Interestingly, multiple nucleoporins are acetylated in yeast (Henriksen et al., 2012), and we found that the acetylation levels of the nuclear basket protein Nup60, and of the central channel components Nup49, Nup53 and Nup57 were reduced in *HOS3-NLS* relative to *hos3*Δ strains (Fig. **7A**), suggesting that these proteins are Hos3 substrates in daughter cells. We next generated point mutants of these nucleoporins in which previously identified acetylated lysines (Henriksen et al., 2012) were changed to asparagine, whose biophysical properties resemble those of acetylated lysine and thus may mimic constitutive acetylation (Ben-Shahar et al., 2008). Consistent with this, lysine-to-asparagine (*KN*) alleles of Nup60, Nup49, Nup53 and Nup57 did not perturb cell growth or Nup60 perinuclear localization, and their combination conferred resistance to the toxic effects of Hos3-NLS over-expression (Fig. **S15** and Table **S5**).

**Figure 7:**
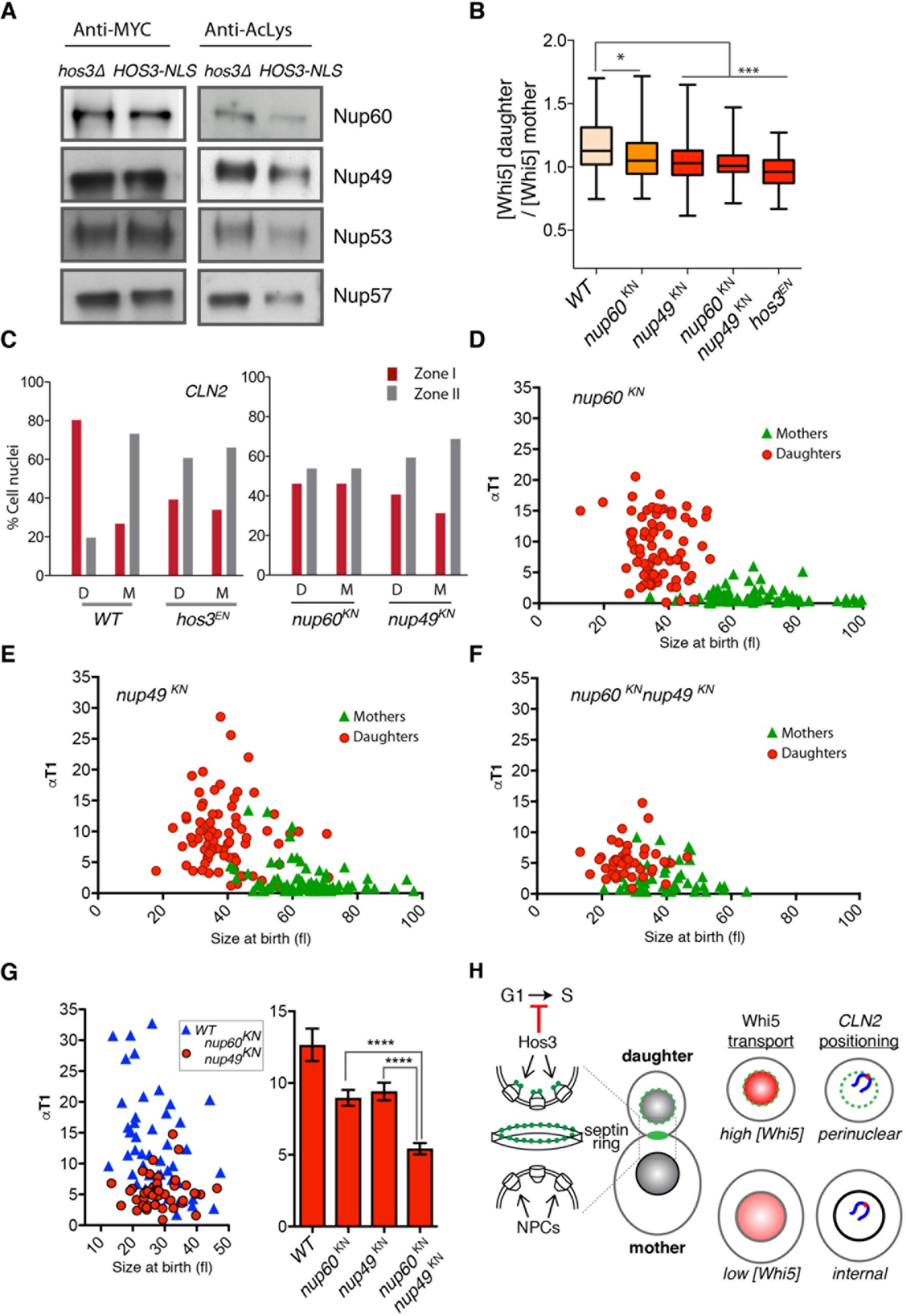
Modification of nucleoporins by Hos3 regulates cell cycle entry. **(A)** Acetylation of nucleoporins in Hos3 mutants. Endogenously tagged proteins were immunoprecipitated from *HOS3-NLS* and *hos3*Δ extracts. Protein levels were assessed with anty-myc antibodies and acetylation levels were assayed with an antibody against acetylated lysines. **(B)** Distribution of Whi5-GFP in wild-type and acetyl-mimic (*KN*) mutants of the indicated nucleoporins. Nuclear fluorescence was measured in mother and daughter cells at the time of birth as in Fig. 5C (> 130 cells / strain; *p* < 0.05 (*) and 0.001 (***), Mann-Whitney test). **(C)** Subnuclear position of *CLN2* (as in Fig. 6D) during the first 24 minutes following cytokinesis, determined by time-lapse imaging in daughter (D) and mother cells (M) of the indicated strains. Images were acquired at 12 min intervals. (N > 120 cells per strain). **(D-F)** Correlation between αT1 and cell size at the time of cytokinesis, for mother and daughter cell pairs in the indicated strains (N>90 cells). **(G)** Correlation between αT1 and cell size at the time of cytokinesis for daughter cells of the indicated strains (left) and αT1 values (mean and SEM) for cells of size at birth 25 - 50 f1 for cells shown in (D-F) (***, *p* < 0.001; Mann-Whitney test) (right). **(H)** Model for Hos3 function in cell cycle entry. Hos3 concentrates at the daughter side of the septin ring at bud neck prior to anaphase. Upon passage of the nucleus through the bud neck, Hos3 is imported in a Mtr10-dependent manner and associates with the nuclear basket of NPCs. Hos3 deacetylates NPCs to affect their functions in daughter cells, leading to perinuclear positioning of the silent *CLN2* locus and to higher accumulation of the Whi5 transcriptional repressor in daughter nuclei, delaying Start in daughter cells.

We next used time-lapse microscopy to determine whether nucleoporin *KN* alleles affect asymmetric Whi5 nuclear concentration and perinuclear tethering of *CLN2* in daughter cells. We found that asymmetric partitioning of Whi5-GFP was mildly affected in *nup60*^*KN*^ cells, and was severely impaired in cells expressing *KN* alleles of central channel nucleoporins, to levels similar to cells lacking Hos3 activity (Fig. **7B** and **S16A**). Moreover, *CLN2* perinuclear tethering was defective in early G1 *hos3*^*EN*^, *nup49*^*KN*^ and *nup60*^*KN*^ daughters (Fig. **7C**). Thus, constitutive acetylation of putative Hos3 substrates at the NPC disrupts partitioning of Whi5 and perinuclear tethering of *CLN2*.

Finally, we examined progression through Start in single and double *KN* mutant strains. This analysis showed a moderate reduction of T1 in single *KN* mutant daughters relative to wild type daughters of similar size (Fig. **7D, E, G** and **S16B**). Remarkably, T1 duration was strongly reduced in *nup49*^*KN*^ *nup60*^*KN*^ double mutant daughters to levels comparable to *hos3* mutants (Fig. **7F-G**). Thus, a combination of these two *KN* alleles mimics the effect of Hos3 inactivation in daughter cell cycle control, suggesting that constitutive acetylation of Nup60 and Nup49 is sufficient to bypass Hos3-mediated inhibition of cell cycle Start in daughter cells.

## Discussion

Here, we have identified a mechanism that establishes differences in NPC acetylation and cell fate between mother and daughter cells in *Saccharomyces cerevisiae*. The Hos3 deacetylase, which localizes to the daughter side of the bud neck during mitosis, associates with NPCs in the daughter cell to direct its cell cycle program. This stands in stark contrast to other known fate-determinant partition mechanisms, which rely on compartmentalization and polarization of the cell cortex. The enrichment of Hos3 at the nuclear periphery in daughter cells was dependent on septin ring integrity, on the septin-associated protein Hsl7, and on passage of the nucleus through the bud neck. Although it is formally possible that local protein degradation contributes to Hos3 localization dynamics, we speculate that NPC association is triggered by the close proximity between the daughter side of the septin ring and NPCs traversing the bud neck.

Daughter-specific inheritance of Hos3 delayed cell cycle commitment of daughter cells. Indeed, deletion of Hos3 shortened G1 duration in daughters but did not affect cell cycle progression of mother cells. Conversely, artificial recruitment of Hos3 to the NPCs of mother cells was sufficient to delay their G1/S transition (Start), indicating that Hos3 acts at the NPCs to inhibit cell cycle entry. Hos3 promotes activity of the spindle orientation checkpoint, a mechanism that prevents exit from mitosis in cells with misoriented spindles (Wang and Collins, 2014). This function is likely associated with localization of Hos3 to the daughter spindle pole. Our findings reveal that an additional, separate pool of Hos3 localizes to the NPC to control cell cycle entry. Furthermore, our data suggest that Hos3 delays Start through mechanisms independent from Ace2, which represses Cln3 transcription in daughter cells (Di Talia et al., 2009; Laabs et al., 2003), and from Rpd3, a type I HDAC that associates with the *CLN2* promoter and favors its repression (Huang et al., 2009).

Our results define two key processes regulated by Hos3 to inhibit the G1/S transition. Firstly, Hos3 is required for enrichment of the Start inhibitor Whi5 in daughter nuclei (Liu et al., 2015; Schmoller et al., 2015). Whereas Hos3 promotes higher concentration of Whi5 in daughter nuclei, it counters the accumulation of karyopherins Kap95 and Msn5 in daughters, which control nuclear levels of Whi5 (Kosugi et al., 2009; Taberner et al., 2009). Since the concentration of Whi5 and the probability to enter the cell cycle are inversely correlated, this provides a mechanism by which Hos3 can control Start in daughter cells. Secondly, we find that Hos3 confines the *CLN2* gene locus to the nuclear periphery in G1 daughter cells, and that perinuclear *CLN2* anchoring delays Start. Hos3 might target *CLN2* to repressive perinuclear domains or to NPCs in daughter cells. Nucleoporins can directly activate gene expression at NPCs in yeast (Casolari et al., 2004; Taddei et al., 2006) or at the nucleoplasm, in metazoans (Capelson et al., 2010b; Kalverda et al., 2010; Liang et al., 2013); however, silencing of subtelomeric genes after association with NPCs has also been reported (Feuerbach et al., 2002; Van de Vosse et al., 2013). In any case, perinuclear silencing of *CLN2* would dampen the positive feedback on Cdk1, further contributing to delaying Start in daughter cells. Interestingly, acetylation by the SAGA complex plays a major role in the recruitment of chromatin to NPCs and the nuclear periphery (Denoth-Lippuner et al., 2014; Dultz et al., 2016). Whether Hos3 counteracts SAGA to control Start in daughter cells remains to be determined.

Our data indicates that Hos3 acts to repress Start by modulating the functions of nuclear pore complexes. Notably, Hos3 physically interacts with the nuclear basket component Nup60, and perinuclear Hos3 deacetylates basket and central channel nucleoporins. Mutation of individual acetylated lysines in either Nup60 or the central channel component Nup49 perturbs Whi5 and *CLN2* asymmetries, and a combination of these mutations entirely disrupts Start inhibition in daughter cells. Based on these observations, we propose that Hos3 inhibits cell cycle entry in daughter cells through deacetylation of the nuclear basket (which predominantly modulates *CLN2* tethering) and of the central channel, which affects both *CLN2* positioning and Whi5 partitioning (Fig. **7H**). Although both Whi5 nuclear levels and *CLN2* gene positioning are Hos3-dependent, these two processes are likely regulated independently of each other. Supporting this possibility, Whi5 is not required for *CLN2* perinuclear anchoring in G1, and *nup60* mutations perturb *CLN2* anchoring but not Whi5 asymmetry. It is important to note that Hos3 may have additional substrates at the NPC that may also play a role in cell cycle regulation. Furthermore, Hos3 may control the nucleo-cytoplasmic shuttling of other factors besides Whi5, and sub-nuclear positioning of other genes besides *CLN2*, which may contribute to the daughter-specific Start delay.

In summary, we reveal that asymmetric partitioning of a lysine deacetylase during mitosis modulates NPC properties in daughter cells to promote two distinct mechanisms underlying their cellular identity. These data reveal that modulation of NPC functions plays a key role in the control of cell cycle progression, and establish deacetylation as a mechanism to regulate NPCs in different cell cycle stages and cell types. The segregation mechanism of Hos3 may be specific to budding yeasts, which have a specialized type of nuclear division. However, differential expression or localization of Hos3-like HDACs might modulate NPC acetylation in other organisms in a cell cycle- or cell type-specific manner. Interestingly, the human homologues of yeast Nup60 and Nup49 (Nup153 and Nup58, respectively) are also acetylated (Choudhary et al., 2009). Moreover, an interplay between NPCs and tissue-specific class-II HDACs has been linked to the regulation of gene expression and positioning in cultured animal cells (Brown et al., 2008; Kehat et al., 2011; Zullo et al., 2012) and Nup153 is involved in ES cell pluripotency through gene silencing (Jacinto et al., 2015). Whether any of these processes is controlled by acetylation of NPCs is not known. Thus, our findings open the possibility that nucleoporin deacetylation might be a conserved mechanism to regulate NPC functions in specific cell types. It will be of interest to determine whether as in budding yeast, NPC deacetylation can affect the establishment of cell identity in multicellular organisms.

## Experimental Procedures

### Strains, cell growth

*S. cerevisiae* strains are derivatives of S288c except when noted (strains; Table S3). Gene deletions and insertions were generated by standard one-step PCR-based methods (Janke et al., 2004) (oligos; Table S4). Site-directed mutagenesis (QuickChange, Agilent Technologies) as used to generate the *HOS3pr-HOS3^EN^-GFP*. KN mutants were generated using CRISPR/Cas9 to replace acetylated lysines [identified in (Henriksen et al., 2012)] with asparagine, and are listed in Table S5. Positive clones were generated and verified by sequencing as described (Laughery et al., 2015). At least two independent clones per strain were analyzed, with identical results. LacO arrays were inserted using a two-step PCR based method (Rohner et al., 2008). LexA or LexA-Yif1 was expressed from pRS425-based plasmids under the control of the *ADH1* promoter. For G1 arrest, cells were grown in YPDA (yeast extract, peptone, dextrose, and adenine) to log phase, synchronized with 15 μg/ml alpha-factor (Sigma-Aldrich) for 2 h and released in fresh YPDA at 30°C. For induction of the *GAL1,10* promoter, cells were grown overnight in YP+Raffinose medium, washed, shifted to YP+Galactose medium for 4 hours, and then plated on YP+Gal plates.

### Fluorescence microscopy

Time-lapse imaging was performed using a spinning-disk confocal microscope (Revolution XD; Andor Technology) with a Plan Apochromat 100x, 1.45 NA objective equipped with a dual-mode electron-modifying charge-coupled device camera (iXon 897 E Dual Mode EM-CCD camera; Andor Technology), temperature-controlled microscopy chamber, a z-stepper and an automated stage. Images were analyzed on 2D maximum projections. For epifluorescence microscopy, a Leica AF 6000 wide-field microscope with an Andor DU-885K-CSO-#VP camera was used. Maximum projections are shown throughout, except in Fig. 1A, where one confocal section is shown for clarity. For tethering of Hos3 to nuclear pores, inducible dimerization of FK506-binding protein (FKBP) and FKBP-rapamycin binding (FRB) domain was used with C-terminally tagged proteins, as described (Gallego et al., 2013). The final strain harbors the *tor1-1* mutation and lacks the endogenous *FPR1* gene rendering growth insensitive to rapamycin. Cells were incubated with 25 μM rapamycin throughout the imaging experiment. Quantification of GFP fusion protein abundance was determined in background-subtracted 2D sum projections of whole-cell Z-stacks, with the nuclear area defined by Nup49-mCherry. The position of GFP-LacI spots was mapped in confocal sections by assigning the GFP spot to one of three nuclear zones of equal area, as described (Hediger et al., 2002). Analysis of G1 duration was performed essentially as described (Di Talia et al., 2007) except that cell size was measured on DIC stacks using ImageJ and a customized version of the BudJ3D plug-in (Ferrezuelo et al., 2012) to calculate cell volume using bright-field images. Growth rates were determined by linear regression of volumetric data over time. Time-lapse series of 4 μm stacks spaced 0.2-0.3 μm were acquired every 2-4 minutes.

### Immunoprecipitation

Cells were grown to log phase or synchronized by Cdc20 depletion, lysed in bead beater with glass beads, and extracts incubated with anti-Myc antibody (05-724; Millipore) coupled to Dynabeads M-280 sheep anti-mouse IgG (11201D) to immuno-precipitate Hos3-myc. For western blotting, anti-Myc (A-14:sc-789, Santa Cruz), anti-GFP (Ref 11814460001, Roche) and anti-acetyl Lys (Cell Signaling, Ref 9681) antibodies were used.

### Author Contributions

Conceptualization, A.K. and M.M.; Methodology, A.K., A.D., Z.S., I.M., P.S., and C.F.; Investigation, A.K., A.D., Z.S., I.M., P.S., T.S. and C.F.; Writing – Original Draft, M.M.; Writing – Review & Editing, A.K. and M.M.; Funding Acquisition, M.B. and M.M.; Supervision, M.M.

## Acknowledgments

We thank Martí Aldea, Jan Skotheim and all members of the Mendoza lab for discussions and sharing reagents, and Yves Barral, Lucas Carey, Ben Lehner and Life Science Editors for comments and manuscript editing. We are grateful to Frederick Cross (Rockefeller University), Josep Clotet (Universtat Internacional de Catalunya), Pedro Carvalho, Vivek Malhotra (both CRG, Barcelona), Susan Gasser (FMI, Basel), Karim Mekhail (University of Toronto) and Andrew Murray (Harvard University) for reagents, the CRG Advanced Light Microscopy Unit (Timo Zimmermann, Raquel Gartía), and Yannick Schwab (EMBL, Heidelberg) for electron microscopy. We acknowledge support the European Research Council (ERC) Starting Grant 2010-St-20091118 to M.M., and the Spanish Ministry of Economy and Competitiveness, ‘Centro de Excelencia Severo Ochoa 2013-2017’, SEV-2012-0208 to the CRG.

The authors declare no conflict of interests.

## Supplementary Information

Extended Experimental Procedures, Figures S1-S16, Tables S1-S3, Movies S1-7

